## Supplementary figures and images for "A Non-Genetic, Cell Cycle Dependent Mechanism of Platinum Resistance in Lung Adenocarcinoma"

### Figure 1 - Supplement 1

#### Figure 1 - Supplement 1 Rajal et al 2021

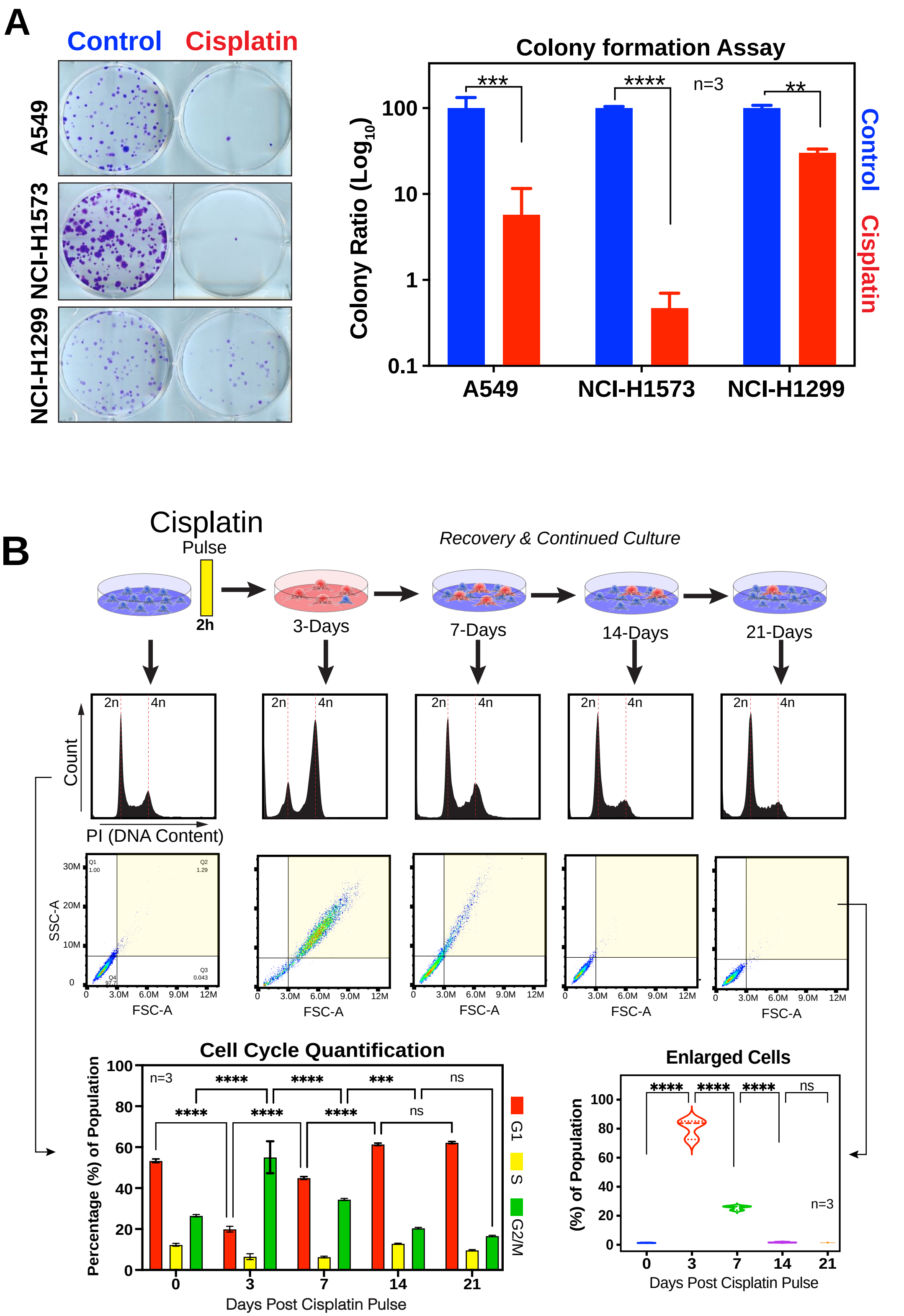

### Figure 2 - Supplement 1

#### Figure 2 -Supplement 1

Rajal et al 2021

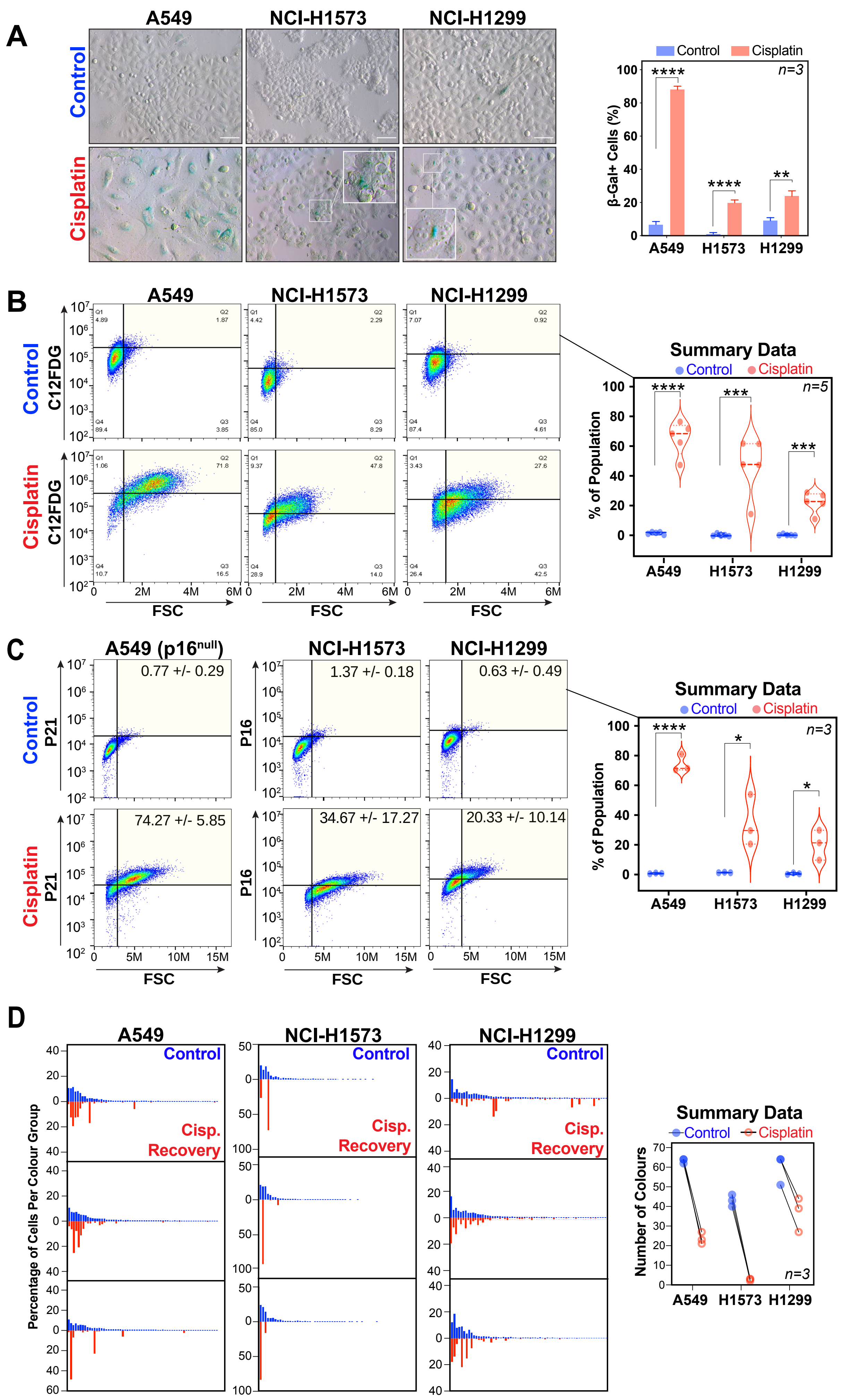

### Figure 3 - Supplement 1

#### Figure 3 -Supplement 1

Rajal et al 2021

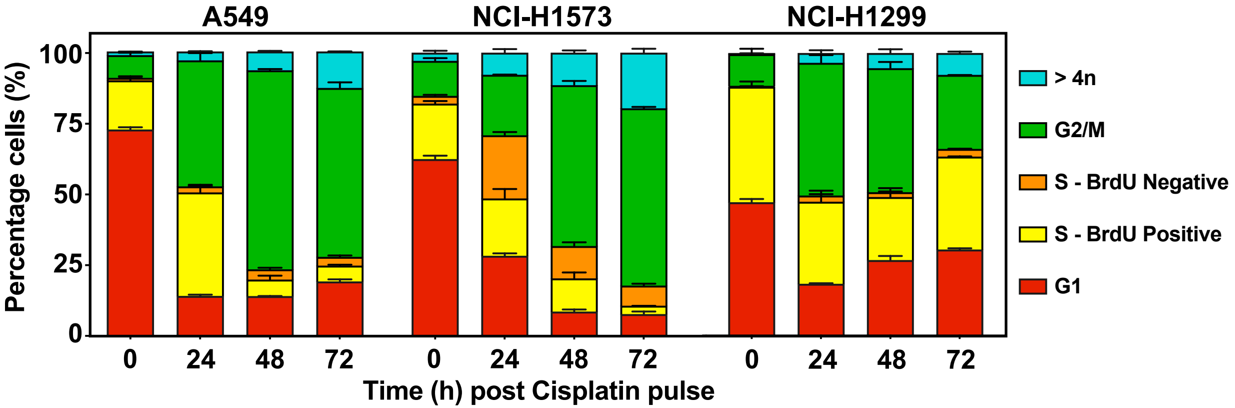

### Figure 4 - Supplement 1

#### Figure 4 - Supplement 1

Rajal et al 2021

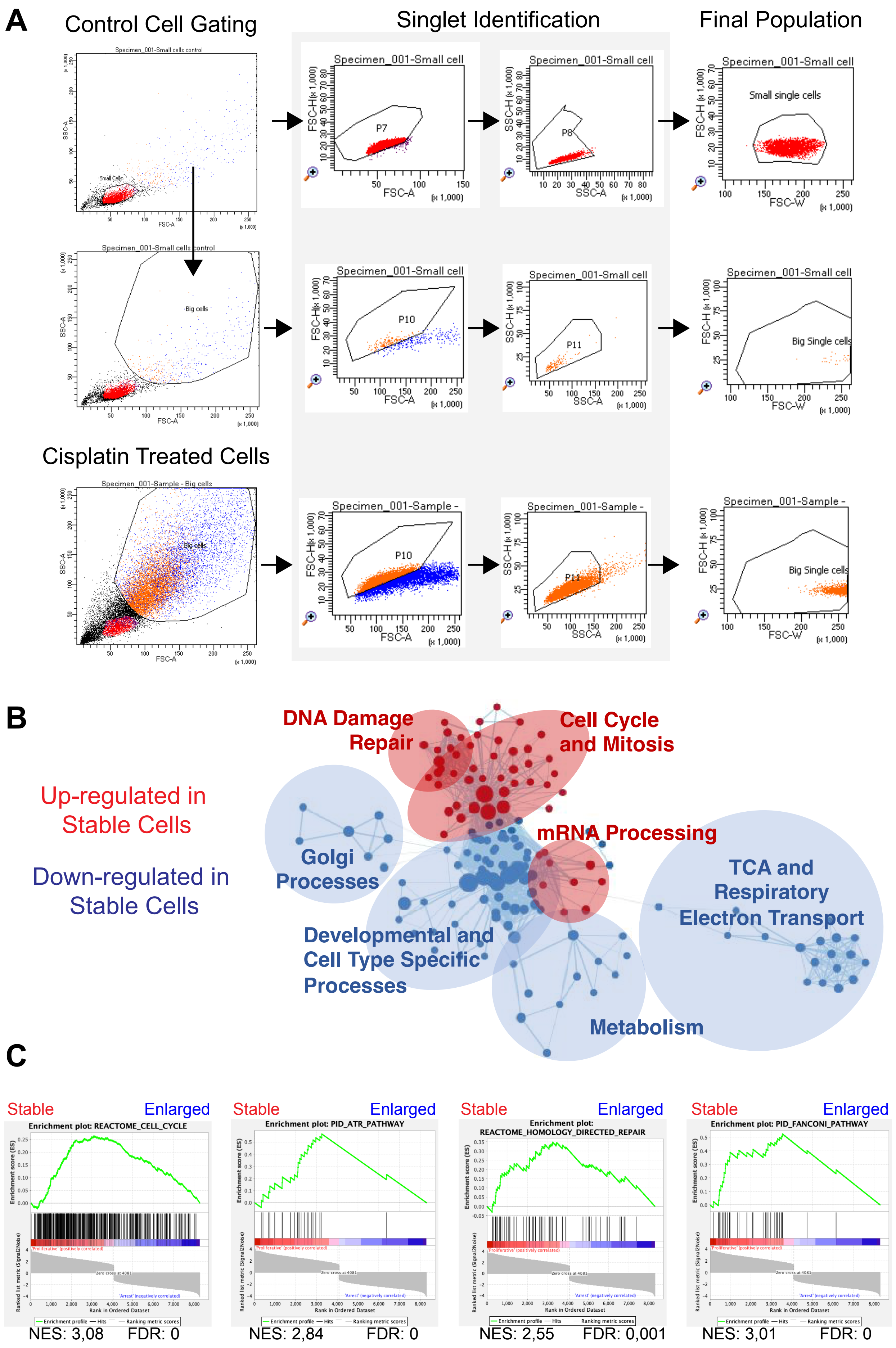

### Figure 5 - Supplement 1

#### Figure 5 -Supplement 1

Rajal et al 2021

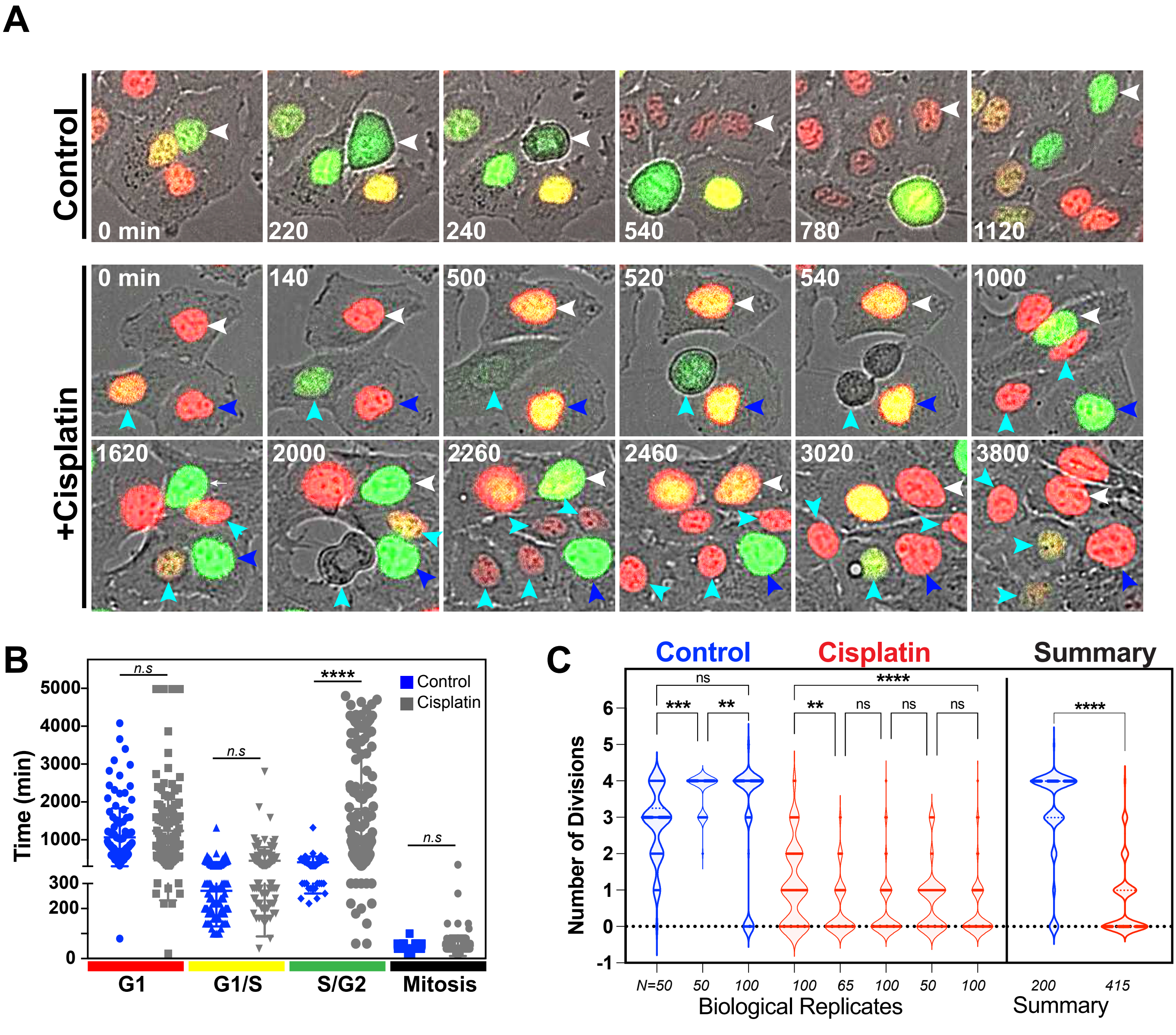

### Figure 5 - Supplement 2

#### Figure 5 -Supplement 2

Rajal et al 2021

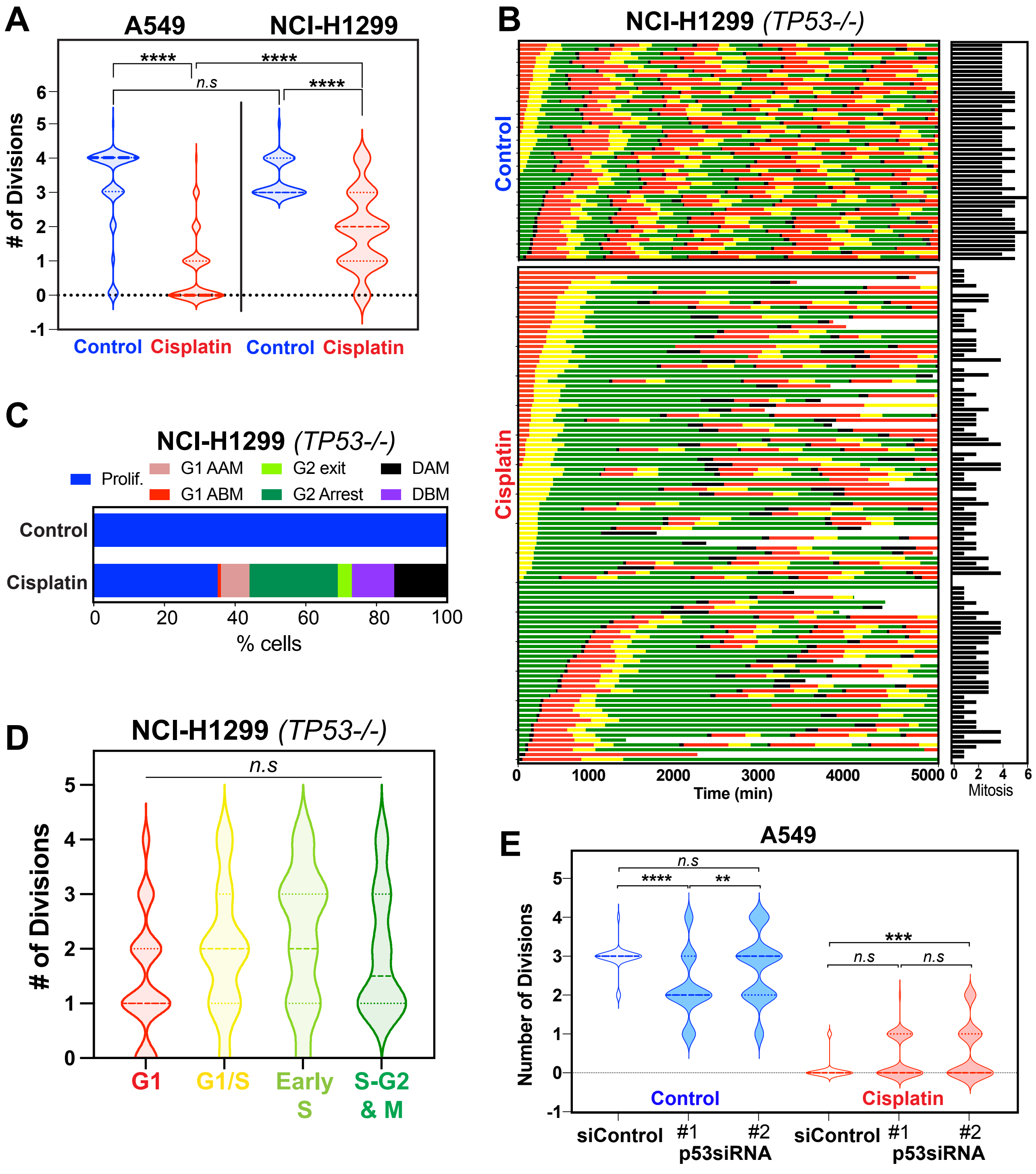

### Figure 5 - Supplement 3

#### Figure 5 -Supplement 3 Rajal et al 2021

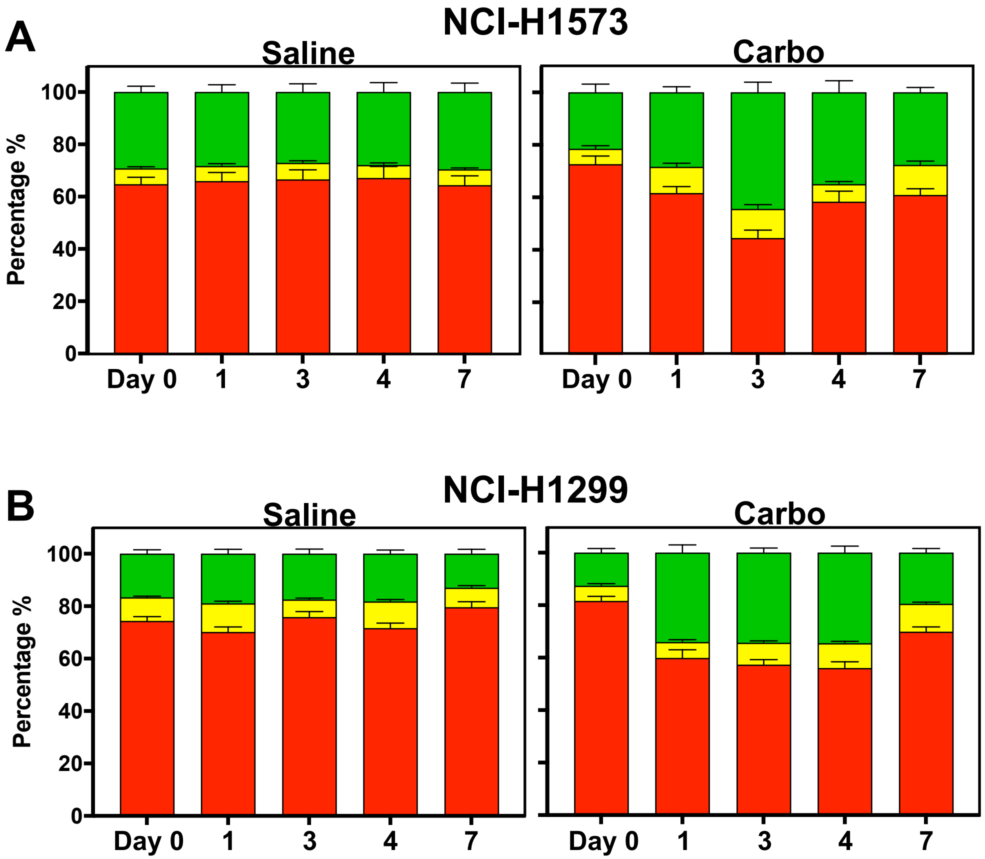

### Figure 7 - Supplement 1

#### Figure 7 - Supplement 1

Rajal et al 2021

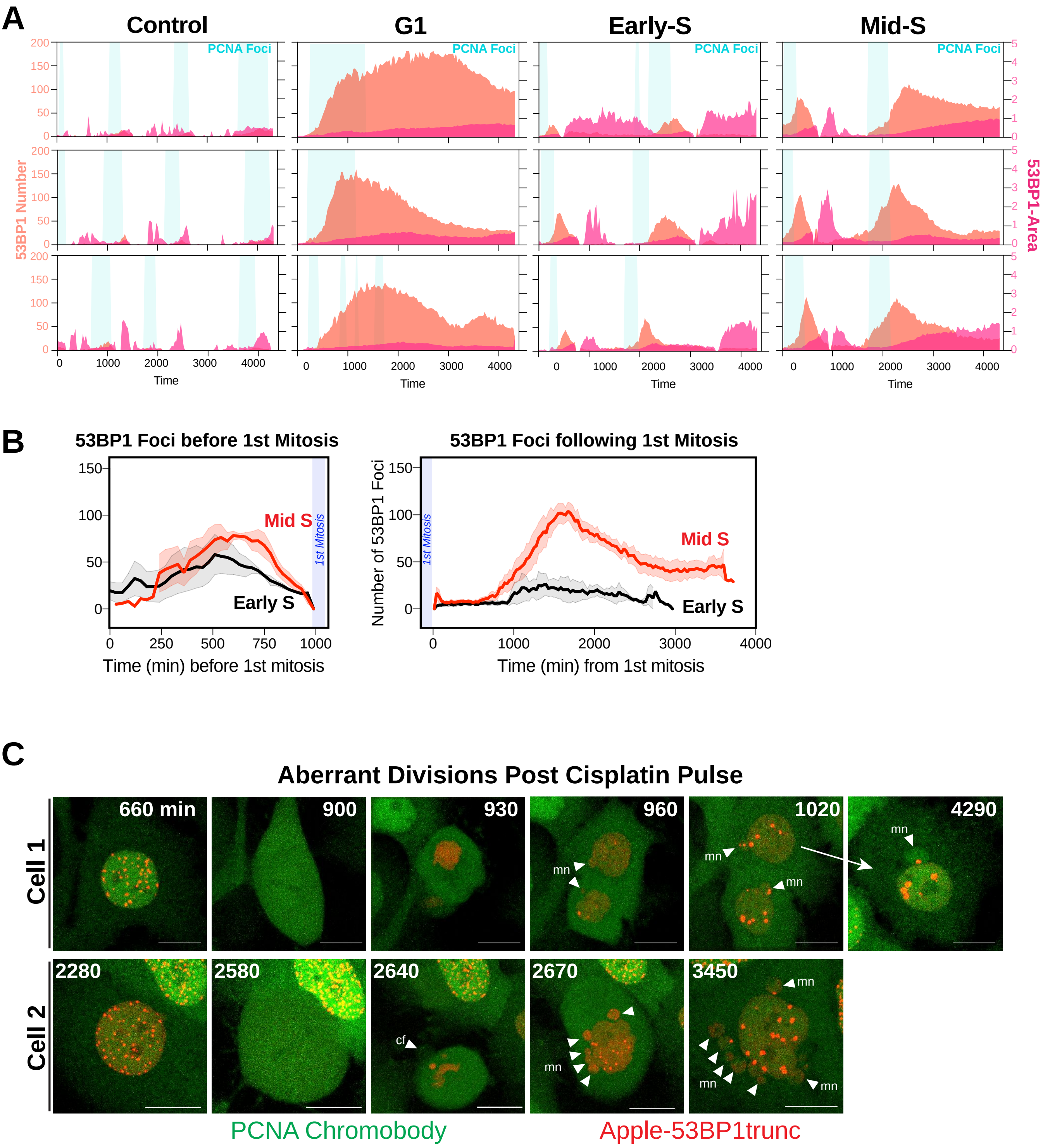

### Figure 8 - Supplement 1

#### Figure 8 Supplement 1

Rajal et al 2021

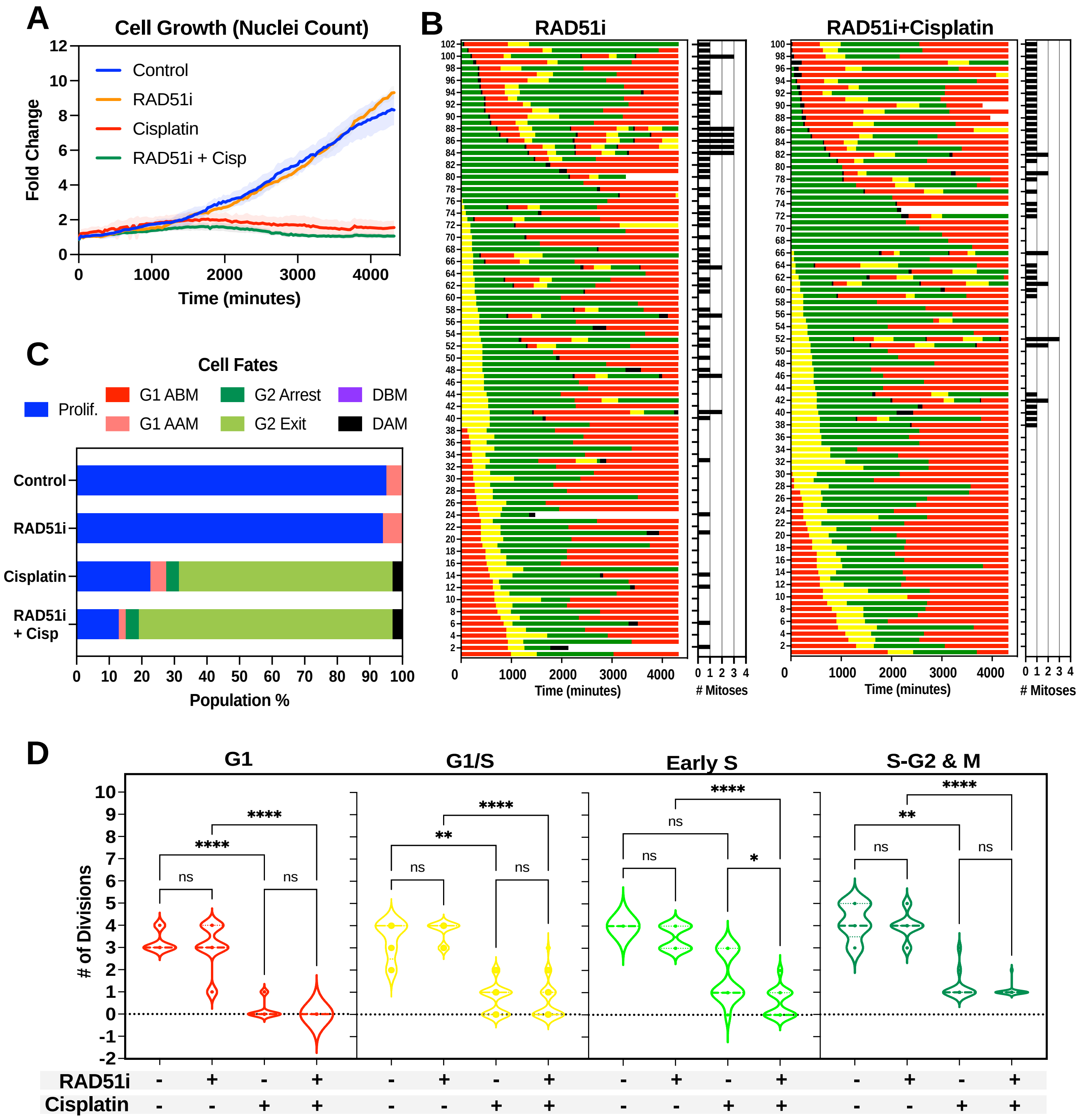
